# Risk of soil-transmitted helminthiasis among agrarian communities of Kogi State, Nigeria: Evaluated in the context of The Soil-Transmitted Helminthiasis Advisory Committee recommendation 2016

**DOI:** 10.1101/663237

**Authors:** Joy T. Anunobi, Ikem C. Okoye, Ifeanyi Oscar N. Aguzie, Yvonne E. Ndukwe, Onyekachi J. Okpasuo

**Affiliations:** Science Laboratory Technology Department, Federal Polytechnic, Idah, Kogi State, Nigeria; Parasitology and Public Health Unit, Department of Zoology and Environmental Biology, University of Nigeria, Nsukka, Enugu State, Nigeria

**Author notes:** Corresponding authors: (IONA), (ICO). **Authors’ contributions** **Conceptualization and Methodology**: Joy T. Anunobi and Ikem C. Okoye **Investigation**: Joy T. Anunobi, Yvonne E. Ndukwe and Onyekachi J. Okpasuo **Data curation, formal analysis & software**: Ifeanyi Oscar N. Aguzie **Writing of first draft**: Joy T. Anunobi and Ifeanyi Oscar N. Aguzie **Supervision:** Ikem C. Okoye **Final draft:** All the authors approved the final draft. **Funding** The study did not receive external funding.

**Keywords:** KAP, socio-economic, soil-transmitted helminths, *Ascaris lumbricoides*, hookworm, *Trichuris trichiura*, neglected tropical diseases, wealth quintiles

## Abstract

Soil-transmitted helminths (STH) have remained a major threat to human especially children in developing countries including Nigeria. Interventions have always been geared towards school-aged children, neglecting preschool-aged children and occupational risk adults. The Soil-Transmitted Helminthiasis Advisory Committee (STHAC) recently suggested incorporating other at-risk groups. In the context of this recommendation, this study assessed the associated risk of STH infection among agrarian communities of Kogi State, Nigeria. A total of 310 individuals of all ages participated in the cross-sectional survey. Stool samples were analyzed using standard Kato-Katz method. A total of 106 (34.2%) individuals were infected with at least one STH. Hookworm was the most prevalent (18.1%); followed by *Ascaris lumbricoides* (16.8%). Worm intensity was generally light. Prevalence of infection was similar between four age groups considered (preschool, school, ‘women of reproductive age’ and older at-risk group). Poor socio-economic status (SES) was a major risk for STH infection. Using a 20-assets based criteria, 68 (23.1%) and 73 (24.7%) of 295 questionnaire respondents were classified into first (poorest) and fifth (richest) wealth quintiles respectively. Risk of infection with STH was 60% significantly lower in the richest wealth quintile compared to the poorest (Prevalence Ratio (PR) = 0.4843, 95% CI = 0.2704 – 0.8678, p = 0.015). Open defecators were more likely to harbour STH than those who did not (PR = 1.7878, 95% CI = 1.2366 – 2.5846, p = 0.00201). Pit latrine and water closet toilet each approximately reduced STH infection by 50% (p < 0.05). Preventive chemotherapy for all age groups, health education and provision of basic amenities especially toilets are needed in order to achieve the goal toward the 2020 target of STH control.

**Author summary:** Soil-transmitted helminths (STHs) are major cause of morbidities globally, especially among children in developing countries such as Nigeria. Present World Health Organization recommended control strategy solely require preventive chemotherapy targeting preschool-aged children (PSAC) and school-aged children (SAC), and the recently included women of reproductive age (WRA). The Soil-Transmitted Helminthiasis Advisory Committee (STHAC) which is saddled with responsibility of evaluating STHs status and providing appropriate recommendations proposed that preventive chemotherapy be extended to other at-risk groups. This study evaluates this and some other recommendations of STHAC 2016 using sections of a state in Nigeria where soil-transmitted helminthiasis is endemic.

Findings from this study supports recommendations for extension of preventive chemotherapy to other at-risk groups apart from PSAC and SAC. It supports WASH (water, sanitation and hygiene) as integral part of STH control. This finding emphasizes the need for health education and change in attitude which could promote tenets of WASH. And very importantly, the study emphasizes the role of poverty in the persistence of STH transmission. It is the belief of the authors that there is the need for improved socio-economic status for sustainable gains of control efforts.

## Introduction

Soil-transmitted helminths (STH) are intestinal worms whose immature stages require a period of incubation in the soil before becoming infective [1]. The most common are *Ascaris lumbricoides, Trichuris trichiura*, the hookworms (*Ancylostoma duodonale* and *Necator americanus*) and *Strongyloides stercoralis* [2]. They are transmitted by eggs present in infected human faeces, which contaminate the soil in areas where sanitation is poor. STH infections are included in the list of the world’s neglected tropical diseases, NTDs [3], and are the most common infections among the poorest and most deprived communities [2]. Transmission of *A. lumbricoides* and *T. trichiura* is primarily through oral-faecal route (usually by ingestion of parasite eggs in faeces), whereas hookworm species and *S. stercoralis* are through active skin penetration of infective larva.

STH infections are among the most common chronic human infections in most regions of the world; with approximately 1.5 billion people infected globally [2]. It accounts for a global burden of 3.3 million of disability-adjusted life years [4]. Approximately 270 million preschool-aged children (PSAC) and 600 million school-age children (SAC) are at risk of infection. They live in areas where transmission of these parasites is intense and are in need of treatment and preventive interventions [2,5].

Intestinal helminthiasis prevalence in Nigeria has remained unchanged since the 1970s [6]. Nigeria has the highest burden and endemicity [3,7]. A larger fraction of those affected are young children of ages 5 to 14 years living in rural areas and urban slums [6–9]. The major contributors to persistence of infections are cultural, socio-economic and environmental factors [6,7]. Unhygienic and common practice of indiscriminate defecation or dumping excrement have persisted in Nigeria [7,10,11]. There has been little success in the introduction of latrines to rural Nigeria [12]. A temporal data available from 1990 through 2015 indicate a marginal decrease in the use of improved sanitation facilities from 38% to 29%, and small increase in open defecation from 24% to 25% [13]. Currently, Nigeria is ranked as one of the nations in the world with the highest number of people practicing open defecation, estimated at over 46 million people [14].

In Nigeria, there are still states where there are limited to no epidemiological information on STH infections; Kogi State is one of those with limited information. Majority of the communities in Ibaji and Igalamela-Odolu Local Government Areas (LGAs) of Kogi State are farming communities where children and adults are fully exposed to risks of helminth infection. In agrarian communities, practices such as walking on bare feet, open defecation, eating of fallen fruits and raw unwashed vegetables with unwashed hands in farmlands could predispose individuals to STH infections [9,15]. Majority of the agrarian communities are warm and moist for most of the year creating a good environment for the parasites to develop all year round.

Therefore the aim of the present study was to ascertain the associated risk factors of STHs among the agrarian communities of Ibaji and Igalamela-Odolu LGAs. Specifically the objectives were to assess, the prevalence and intensity of soil-transmitted helminthiasis; proportion of individuals with low, moderate and high worm intensity; age-related distribution of STHs infection; socio-economic status as risk for STHs infection; and risks associated with knowledge, attitude an practices (KAP). These objectives were pursued in view of the year 2016 recommendations of The Soil-Transmitted Helminthiasis Advisory Committee (STHAC) in relation to the revision of the 2012 Strategic Plan for STH control [16,17]. The prevalence and intensity of soil-transmitted helminthiasis in the LGAs were assessed in view of WHO 2020 milestones number 3 that all countries requiring PC for STH have less than 1% prevalence of moderate to high intensity [16,17]; though this study was localized to agrarian communities of Kogi State as arbitrary ‘sentinels sites’ (milestone number 2). Kogi State was included in the recent epidemiological mapping of schistosomiasis and soil-transmitted helminthiasis by the Federal Government of Nigeria [18]; but the mapping was only school based, leaving out vulnerable adults. The WHO 2012 Strategic Plan for STH control recommends preventive chemotherapy for SAC and PSAC, and has now included deworming of women of reproductive age. STHAC and some other authors have suggested inclusion of other at risk groups [17]. In order to evaluate the relevance of this recommendation, considering the additional cost incurred from additional drug provision, this study partitioned our study population into four age groups: PSAC (0 – 4yr), SAC (5 – 18 yrs, though 14yrs was not used as the upper limit), ‘women of reproductive age and men of same age bracket’ (19 – 45 yrs), and older at risk group (> 45yrs). This enabled comparison of STHs infections among the age groups. It is believed that where prevalence of STH infection is similar between the age groups, it would support a need for inclusion of all age group in preventive chemotherapy interventions. The choice of agrarian community was due to the persistent risk of infection in such settings, which may justify need for localized mass drug administration (MDA) strategies. Though current WHO STH control strategy depend solely on preventive chemotherapy, the role of socio-economic status and KAP cannot be ignored. The authors believed that the relevance of STHAC re-affirmation of the importance of water, sanitation and hygiene (WASH) in control of STH could be assessed in relation to socio-economic status and KAP. Therefore, socio-economic status of the study population was assessed using asset-based approach, which involved classification of individuals into wealth quintiles in relation to possession of arbitrary but systematically chosen assets.

Findings from this study supports STHAC recommendations for extension of preventive chemotherapy to other at-risk groups, as prevalence of STH infection was not different between the four age groups considered. It similarly supports intergradation of WASH into STH control effort. The finding emphasizes the need for health education and change in attitude which could promote tenets of WASH. And very importantly, the study emphasizes the role of poverty in the persistence of STH transmission. There is the need for improved socio-economic status for sustainable gains of control efforts.

## Methods

### Study design

The study employed a household-based, explorative, cross-sectional study design.

### Study area

The study area comprises Ibaji and Igalamela-Odolu LGAs of Kogi State, Nigeria. In Ibaji LGA, six communities were sampled. They are Ejule-Ojebe, Onyedega, Aneke, Inyano, Ichalla-Ajode and Otiaka. Five communities were sampled in Igalamela-Odolu LGA. They are Ogbagba, Okpachala, Ofuloko, Ogbogbo and Aikpele.

Ibaji LGA is located on the Eastern flank of Kogi State. The north easterly line of equal latitude and longitude passes through the LGA. Ibaji LGA has an area of 1,377 km^2^ and a population of 128,129 [19]. Geographically, it is located between longitudes 6°45’E and 7°00’E and latitudes 6°45’ and 7°00’N. Its inhabitants are primarily Igala speaking tribe [20]. High volume of rainfall and warm temperature result in lush vegetation and good conditions for cultivation of food crops such as rice, vegetables, yam, cassava, sorghum, maize, millet, cowpea, and groundnut which predominate the agricultural practice in the LGA. The major occupation is crop farming with some of them taking to fishing activities to supplement their meals and earn income for themselves. Only few of them are civil servants. The area suffers from poor infrastructural facilities such as absence of good road networks, electricity, pipe-borne water, health centres and communication network [21]; and houses in these communities are mainly made of mud walls and thatched roofs.

Igalamela-Odolu LGA has its headquarters in the town of Ajaka (7°10′16″N and 6°49′35″E). The northeasterly line of equal latitude and longitude passes through the LGA. It has an area of 2,175 km² and a population of 148,020 [19]. Majority of the communities in the LGA are made up of crop farmers while few others are civil servants. Basic amenities such as pipe-borne water, electricity, toilets and accessible roads are lacking in most rural communities in the LGA.

### Study Participants and study size

Communities sampled were selected randomly. The study population consisted of both male and female adults and children who are resident in the selected communities.

Sample size was estimated using the sample size estimation formula by Yamane [22].

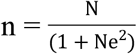

Where, n = sample size, N = total number of study population (based on 2006 census). e = probability level (P = 0.05). The estimated sample size was 399. Though 310 people consented to the study.

Within each chosen community, selection of the first households was by a systematic random sampling. Thereafter alternate households were enlisted. Within each enlisted household, all members were sampled to avoid the usual displeasure of those omitted [23]. Sample collections took place between February and April 2017, though the study duration was between August 2016 and May 2017.

### Ethical Approval

The study protocol was approved by the Ethical Committee of the Kogi State Ministry of Health, Lokoja, Kogi State. The ethical approval was assigned the reference number MOH/KGS/1376/1/79.

### Advocacy and community sensitization

Prior to commencement of the study, advocacy visits were made to designated communities. The traditional rulers of each of the LGAs were visited and briefed on the study objectives. Written permissions were obtained from each of the traditional rulers indicating their consent on behalf of the entire communities. The community and household heads were also well-briefed on the objectives of the study. Verbal permissions were obtained from both the community heads and heads of each household visited. Each member of a household was given a consent form on which interest to participate in the study was indicated. Consent for children was obtained from their parents/care-givers.

### Stool Sample collection and processing

Fresh stool samples for helminth screening were collected from each of the subjects into a coded, dry, leak proof and sterilized sample container, each participant having been briefed to ensure that no urine, water, soil or other contaminants entered the container. The samples were received early in the morning, recorded appropriately and processed immediately. Samples which were not processed in the field were kept refrigerated using a portable cooler and later transported the same day to the laboratory for processing within a maximum of three hours.

Stool specimen was processed using the Kato-Katz technique [24]. Kato-Katz stool examination kit (Vestergaard Fradson, Switzerland) was used. Stool specimen was processed within three hours of collection and examined microscopically within one hour of preparation to avoid over clearance of hookworm eggs [25]. The presence of STH parasites eggs in any of the stool samples was noted as positive. Microscopic examination was repeated days after the slide preparation for the confirmation of the presence of *Trichuris trichiura* and *Ascaris lumbricoides* eggs as a quality control procedure.

### Assessment of demographic and socio-economic status

#### Use of questionnaires

A well-structured, pre-tested questionnaire was administered to each participant. The questionnaire was interview-based since most participants had no formal education and could not read nor write. However, those that could read and write provided the information at their convenience. Responses for 0 – 4 year old participants comprising 18.6% (n = 55) of the study population were provided by their parents/guardians.

The questionnaire was sectioned into nine: (i.) basic information, (ii.) perception and knowledge about intestinal helminths, (iii.) community and household characteristics, (iv.) hygiene practices, (v.) farm/home activities, (vi.) use of shoes or sandals, (vii.) use of health services, (viii.) history of deworming, and (ix.) possession of domestic animals.

### Visual inspection of compounds

Complementary information to questionnaire responses was obtained by visual inspections. Inspection of compounds was made and the sanitary condition and presence or absence of sewage disposal system noted. Wearing of shoes at home was assessed by visual observation. Water supply for drinking and washing of hands after using the toilets or urinals was noted. Children playing on sand were noted as well as those walking around the compound without shoes.

### Quantitative variables

Prevalence of infection was computed as proportion of individuals infected to total number examined and presented in percentages. Mean intensity was obtained based on number of eggs per gramme of faeces (epg) of infected individual and categorized into light, moderate and heavy as established by WHO [26]. The number of eggs obtained was multiplied by 24 which is the standard for the Kato-Katz technique. Mean intensity was obtained as average number of parasite eggs per infected individuals. Infection status and questionnaire variables were dummy-coded for statistical analysis purpose. All 310 participants responded to questionnaire. Incomplete response led to exclusion of 15 (4.8%) questionnaires, leaving 295. Prevalence and infection intensity report on the 310 participants was provided while socio-economic status and risk estimates used the 295 properly completed questionnaires.

### Data analysis

Prevalence of infection was compared using chi-square analysis or Fisher’s exact test. Asset-based approach was used to group study participants into different socio-economic status [27,28]. Possession of different assets considered indicators of socio-economic status (SES) was used to develop a single composite variable SES. Twenty assets were used (S1 Table). Principal component analysis (PCA) extracted the key components. Seven principal components were extracted and the first component which explained 24.9% of total variation served as SES; its factor score loading helped group individuals into quintiles [27](S1 Table). Quintile 1 (Q1) represented the poorest who possessed assets associated with lowest extreme of SES, or who did not possess most assets associated with wealth. Quintile 2 (Q2) were poor, Q3 were average (neither poor nor rich), Q4 were rich while Q5 were the richest. The five quintiles were dummy-coded and used for further analysis. Some assets were either combined or excluded; possession of tiled floor was combined with cement floor and both retained as “cement floor” because only 8 persons used tiles. Possession of dog, pig and cow were removed from the PCA because of their scarcity in the population. Possession of these assets and other attributes were compared between the different SES by chi-square analysis. Risk of infection associated with SES, knowledge and perception, attitude and practices are presented as prevalence ratio (PR). Poisson regression model with robust variance was fitted to estimate the PR, 95% confidence interval and associated p-values [29]. Data was analyzed using RStudio [30] and SPSS version 20.0 (IBM Corporation, Armonk, USA). Level of significance was set at 95% probability level.

## Results

### Prevalence and intensity of soil-transmitted helminthiasis in Ibaji and Igalamela-Odolu LGAs

Prevalence of STH in Ibaji and Igalamela-Odolu LGAs was 34.2%, 106 out of 310 persons examined were infected with at least one of the STH, *Ascaris lumbricoides*, hookworm and *Trichuris trichiura*. Hookworm was the most prevalent (18.1%) and *T. trichiura* the least (5.2%) (Table 1). Difference in prevalence of the three STH was significant (χ^2^ = 27.097, p < 0.0001). STH infections were recorded in the two LGAs. Out of 167 and 143 participants, 58 (34.7%) and 48 (33.6%) respectively were infected in Ibaji and Igalamela-Odolu LGAs; the difference was not significant (χ^2^ = 0.046, p = 0.829). Prevalence of STH infection collectively and specifically was similar between the four age groups considered, namely PSAC (0 – 4yr), SAC (5 – 18 yrs), ‘women of reproductive age and men in similar age bracket’ (19 – 45 yrs), and older at-risk groups (> 45 yrs; Fig 1).

**Table 1.** Overall prevalence of soil-transmitted helminths. (N = 310)

|  | No. Infected (%) | Egg count (epg) |  |
| --- | --- | --- | --- |
| STH | 106 (34.2) | Intensity (Mean ± SD) | Min. – Max. |
| *Ascaris lumbricoides* | 52 (16.8) | $216.92 \pm 129.81$ | 48 – 600 |
| Hookworm | 56 (18.1) | $373.71 \pm 318.10$ | 48 – 2232 |
| *Trichuris trichiura* | 16 (5.2) | $156.00 \pm 143.73$ | 48 – 576 |

N = number examined. Epg = egg per gram. SD = standard deviation

**Fig 1.**
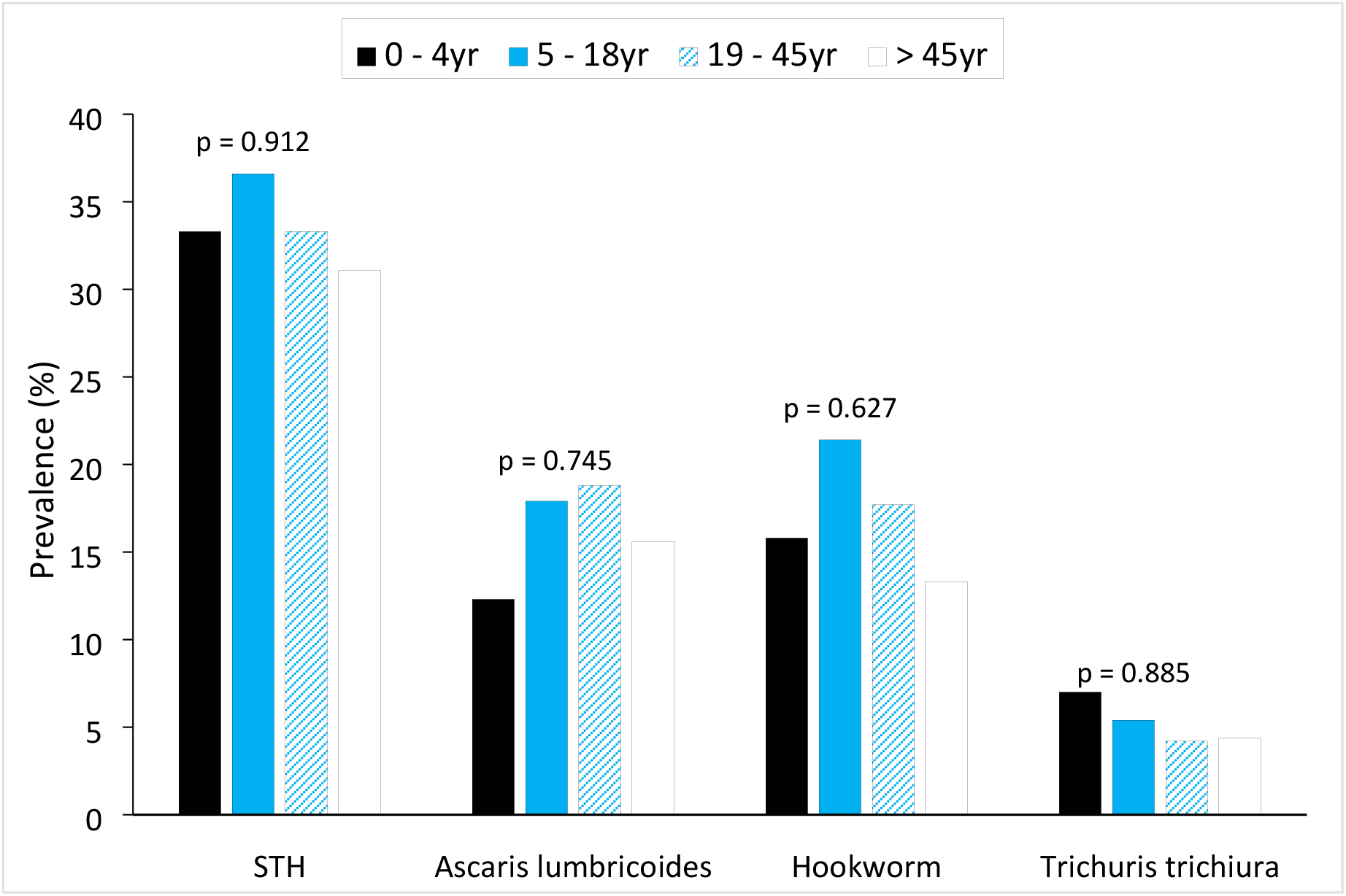
Prevalence of soil-transmitted helminths by age groups.

Mean egg count for hookworm was highest (373.71 ± 318.10 epg, range 48 – 2232) followed by *A. lumbricoides* (216.92 ± 129.81 epg, range 48 – 600) (Table 1). In Fig 2 intensity of the three STH based on WHO [26] standard categories is shown. Intensities of the three helminths were largely light. Collectively, prevalence of moderate and heavy intensity of STH infection was < 1%, though moderate intensity of infection only occurred for hookworm. Moderate intensity of hookworm infection only occurred in 1.8% of individuals infected.

**Fig 2.**
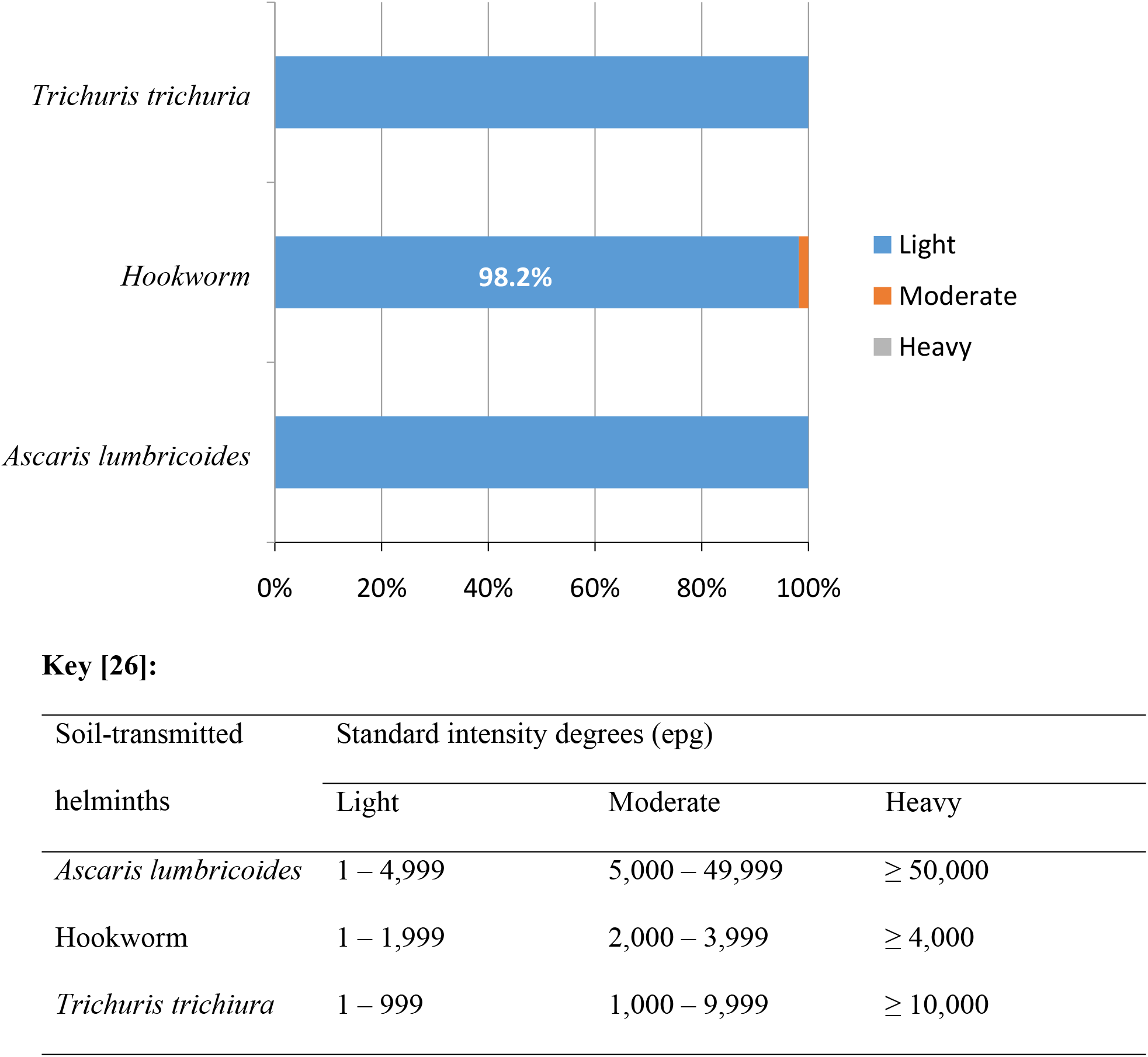
Intensity of soil transmitted helminths in agrarian communities of Ibaji and Igalamela-Odolu Local Government Area, Kogi State.

### Demographics and socio-economic status of questionnaire respondents

All 310 participants responded to questionnaire. Incomplete response led to exclusion of 15 (4.8%) questionnaires, leaving 295. Demographics and SES of respondents are summarized in Tables 2 and 3. Ichalla-Ajode (24.4%) and Ogbagba (4.7%) had the highest and least number of respondents respectively. Majority of the respondents were students (38.6%), farmers (36.9%) and PSAC (21.4%; Table 2). Majority of the participants lived in houses made of mud (50.5%) and cement (46.8%) floors (Table 3). Over 41% of respondents had electricity and 51.9% had pipe-borne water.

**Table 2:** Overall demographics of questionnaire respondents of agrarian communities in Ibaji and Igalamela-Odolu Local Government Areas, Kogi State. (n = 295)

| Characteristics | Frequency (%) |
| --- | --- |
| **LGA** |  |
| Ibaji | 154 (52.2) |
| Igalamela-Odolu | 141 (47.8) |
| **Community** |  |
| Onyedega | 19 (6.4) |
| Ejule | 15 (5.1) |
| Inyano | 15 (5.1) |
| Aneke | 15 (5.1) |
| Ichalla-Ajode | 72 (24.4) |
| Otiaka | 18 (6.1) |
| Ogbagba | 14 (4.7) |
| Okpachala | 28 (9.5) |
| Ofuloko | 18 (6.1) |
| Ogbogbo | 53 (18.0) |
| Aikpele | 28 (9.5) |
| **Sex** |  |
| Male | 153 (51.9) |
| Female | 142 (48.1) |
| **Age (year)** |  |
| 0 – 4 | 55 (18.6) |
| 5 – 18 | 107 (36.3) |
| 19 – 45 | 89 (30.2) |
| > 45 | 44 (14.9) |
| **Marital Status** |  |
| Single | 173 (58.6) |
| Married | 122 (41.4) |
| **Occupation** |  |
| Pre-school | 63 (21.4) |
| Student | 114 (38.6) |
| Farming | 109 (36.9) |
| Fishing | 1 (0.3) |
| Civil Servant | 6 (2.0) |
| Others | 2 (0.7) |

n = number of respondents

**Table 3.**
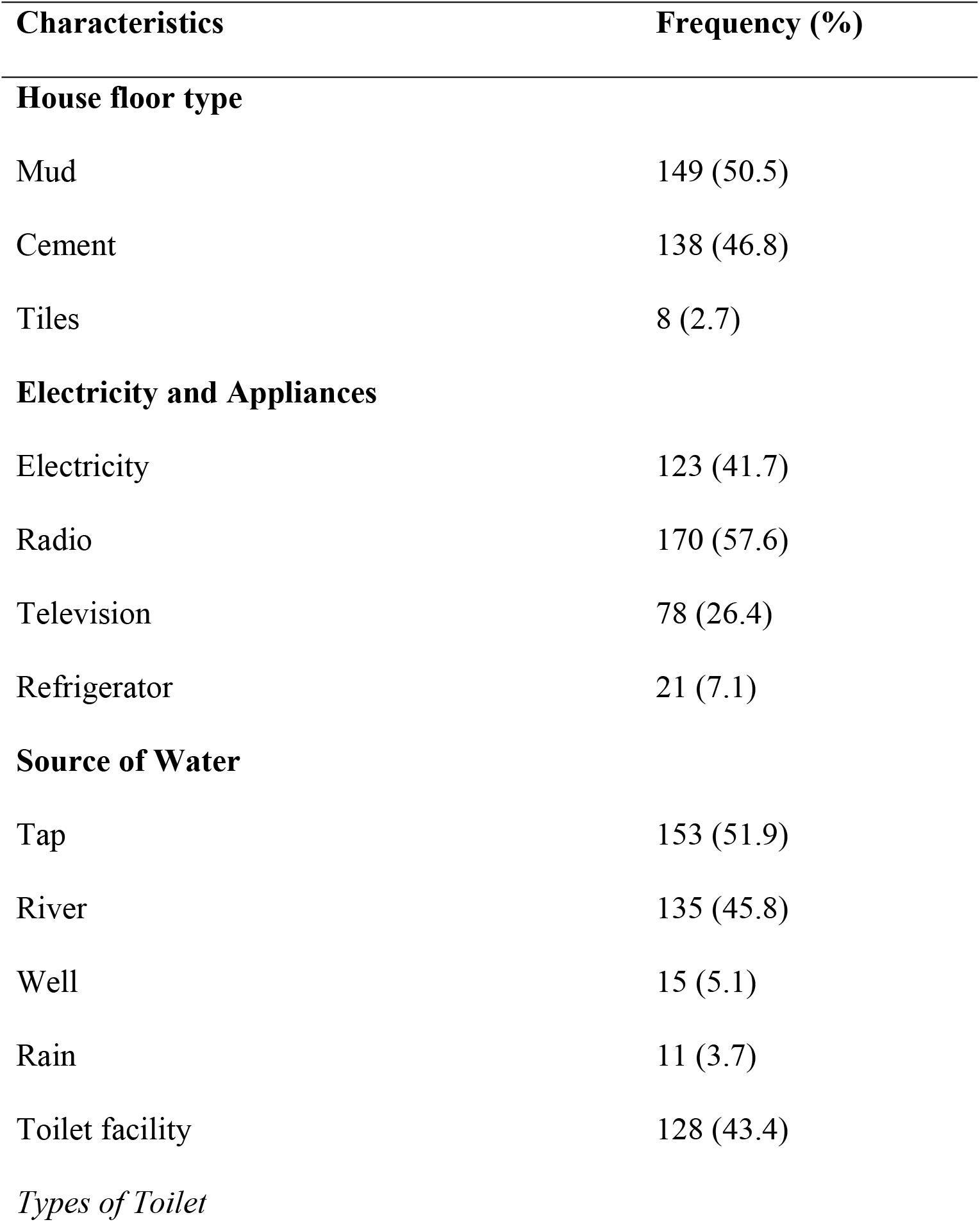
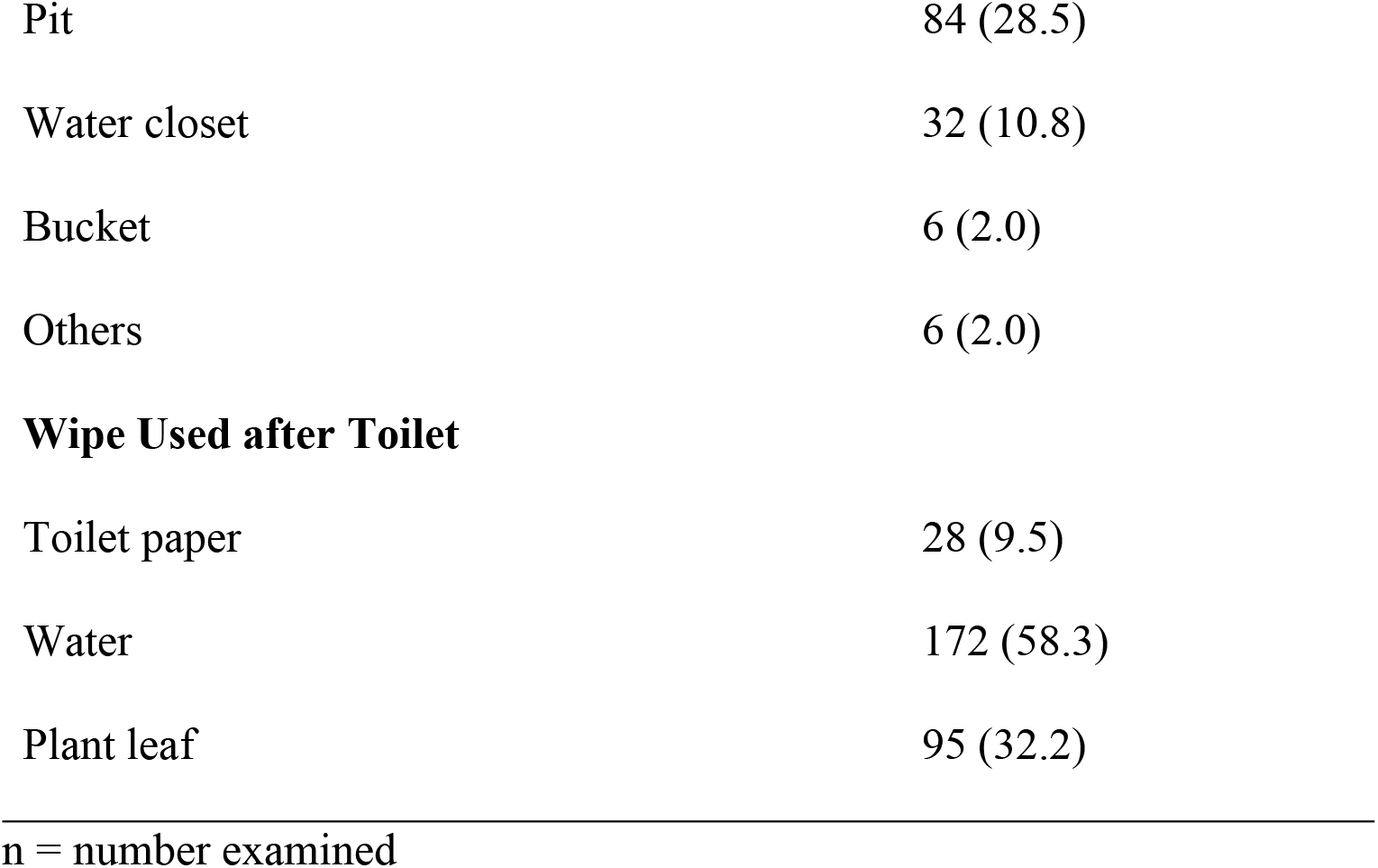
Socio-economic status of questionnaire respondents of agrarian communities in Ibaji and Igalamela-Odolu Local Government Areas, Kogi State. (n = 295).

| Characteristics | Frequency (%) |
| --- | --- |
| **House floor type** |  |
| Mud | 149 (50.5) |
| Cement | 138 (46.8) |
| Tiles | 8 (2.7) |
| **Electricity and Appliances** |  |
| Electricity | 123 (41.7) |
| Radio | 170 (57.6) |
| Television | 78 (26.4) |
| Refrigerator | 21 (7.1) |
| **Source of Water** |  |
| Tap | 153 (51.9) |
| River | 135 (45.8) |
| Well | 15 (5.1) |
| Rain | 11 (3.7) |
| Toilet facility | 128 (43.4) |
| *Types of Toilet* |  |
| Pit | 84 (28.5) |
| Water closet | 32 (10.8) |
| Bucket | 6 (2.0) |
| Others | 6 (2.0) |
| **Wipe Used after Toilet** |  |
| Toilet paper | 28 (9.5) |
| Water | 172 (58.3) |
| Plant leaf | 95 (32.2) |

n = number examined

### Knowledge, perception, attitude and practices

Summary of knowledge, attitude and practices of the questionnaire respondent are presented in Tables 4. Almost half (48.8%) of the respondents think they could get infected by worms but only 8.8% know how they could get infected; and only 16.6% try to prevent infection. A high proportion of respondents defecate in the open (59.3%). Though 64.7% eat food with bare hands (without cutleries), 88.5% washed their hands after using the toilet.

**Table 4.**
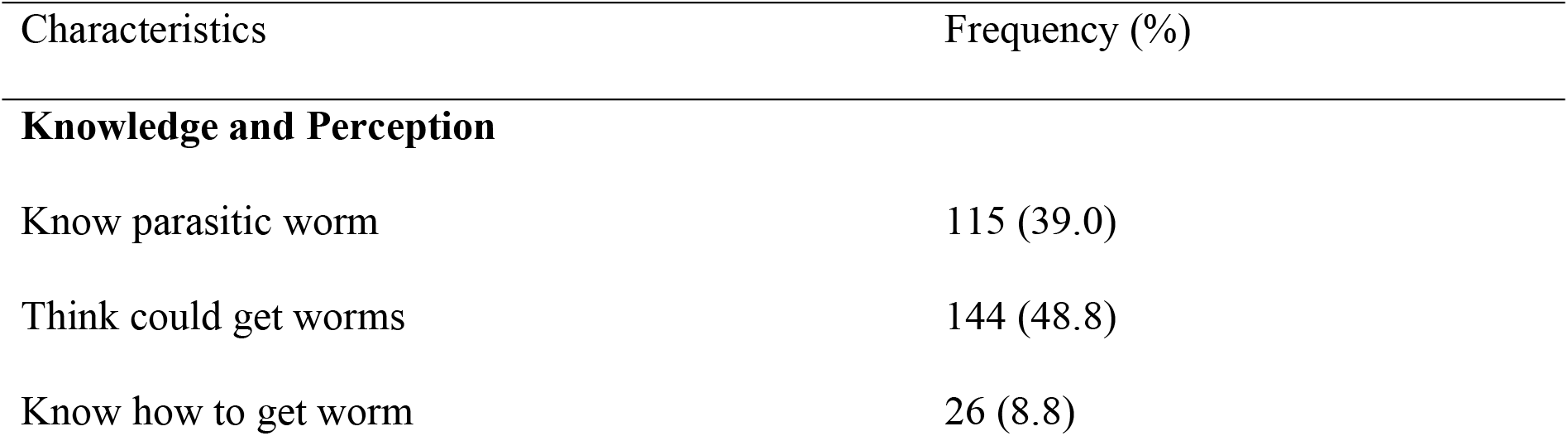
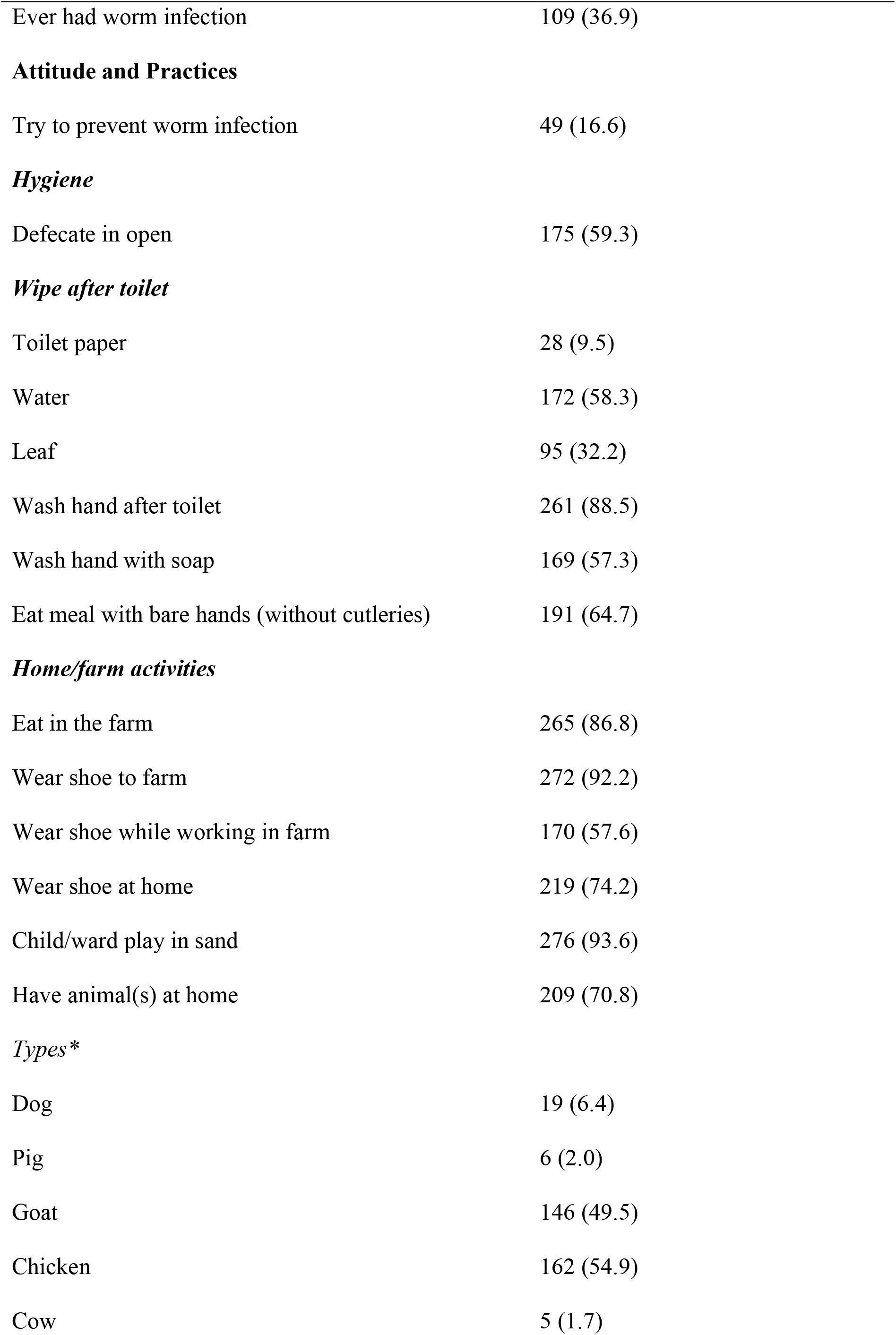
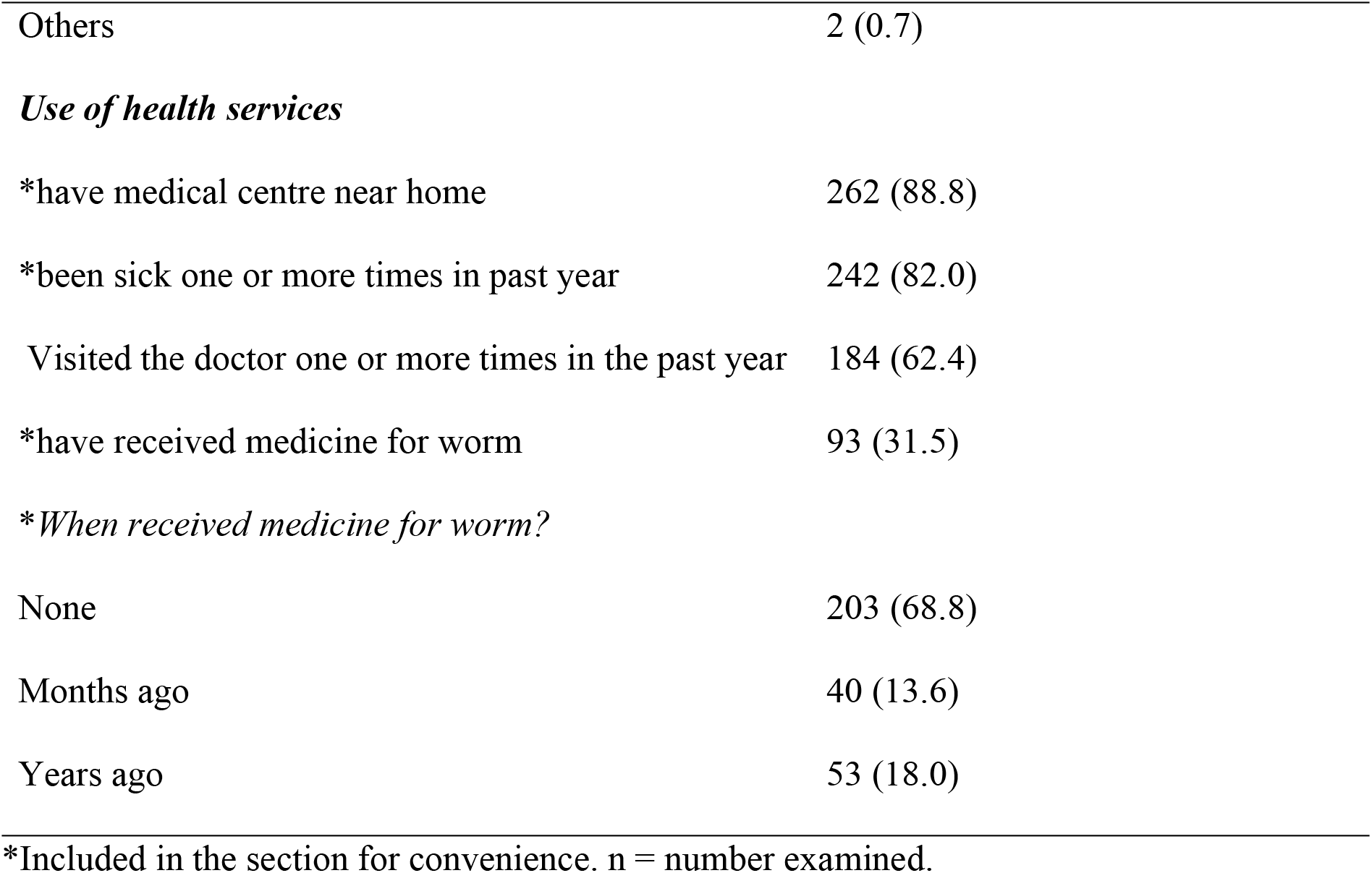
Knowledge, perception, attitude and practice of questionnaire respondents in agrarian communities of Ibaji and Igalamela-Odolu Local Government Area, Kogi State. N = 295.

| Characteristics | Frequency (%) |
| --- | --- |
| **Knowledge and Perception** |  |
| Know parasitic worm | 115 (39.0) |
| Think could get worms | 144 (48.8) |
| Know how to get worm | 26 (8.8) |
| Ever had worm infection | 109 (36.9) |
| **Attitude and Practices** |  |
| Try to prevent worm infection | 49 (16.6) |
| ***Hygiene*** |  |
| Defecate in open | 175 (59.3) |
| ***Wipe after toilet*** |  |
| Toilet paper | 28 (9.5) |
| Water | 172 (58.3) |
| Leaf | 95 (32.2) |
| Wash hand after toilet | 261 (88.5) |
| Wash hand with soap | 169 (57.3) |
| Eat meal with bare hands (without cutleries) | 191 (64.7) |
| ***Home/farm activities*** |  |
| Eat in the farm | 265 (86.8) |
| Wear shoe to farm | 272 (92.2) |
| Wear shoe while working in farm | 170 (57.6) |
| Wear shoe at home | 219 (74.2) |
| Child/ward play in sand | 276 (93.6) |
| Have animal(s) at home | 209 (70.8) |
| ***Types\**** |  |
| Dog | 19 (6.4) |
| Pig | 6 (2.0) |
| Goat | 146 (49.5) |
| Chicken | 162 (54.9) |
| Cow | 5 (1.7) |
| Others | 2 (0.7) |
| *Use of health services* |  |
| *have medical centre near home | 262 (88.8) |
| *been sick one or more times in past year | 242 (82.0) |
| Visited the doctor one or more times in the past year | 184 (62.4) |
| *have received medicine for worm | 93 (31.5) |
| *\*When received medicine for worm?* |  |
| None | 203 (68.8) |
| Months ago | 40 (13.6) |
| Years ago | 53 (18.0) |

\*Included in the section for convenience. n = number examined.

**Table 5.**
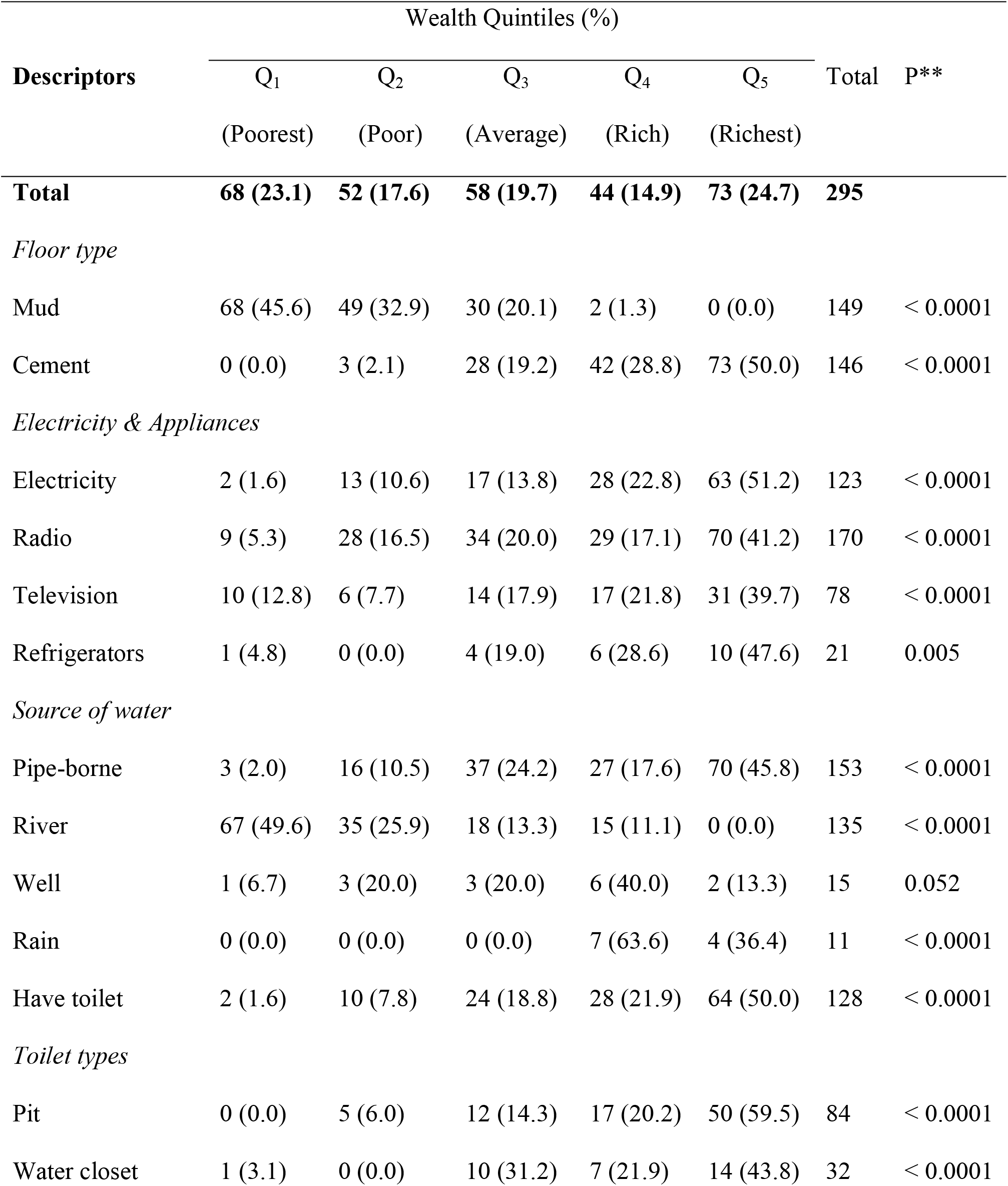
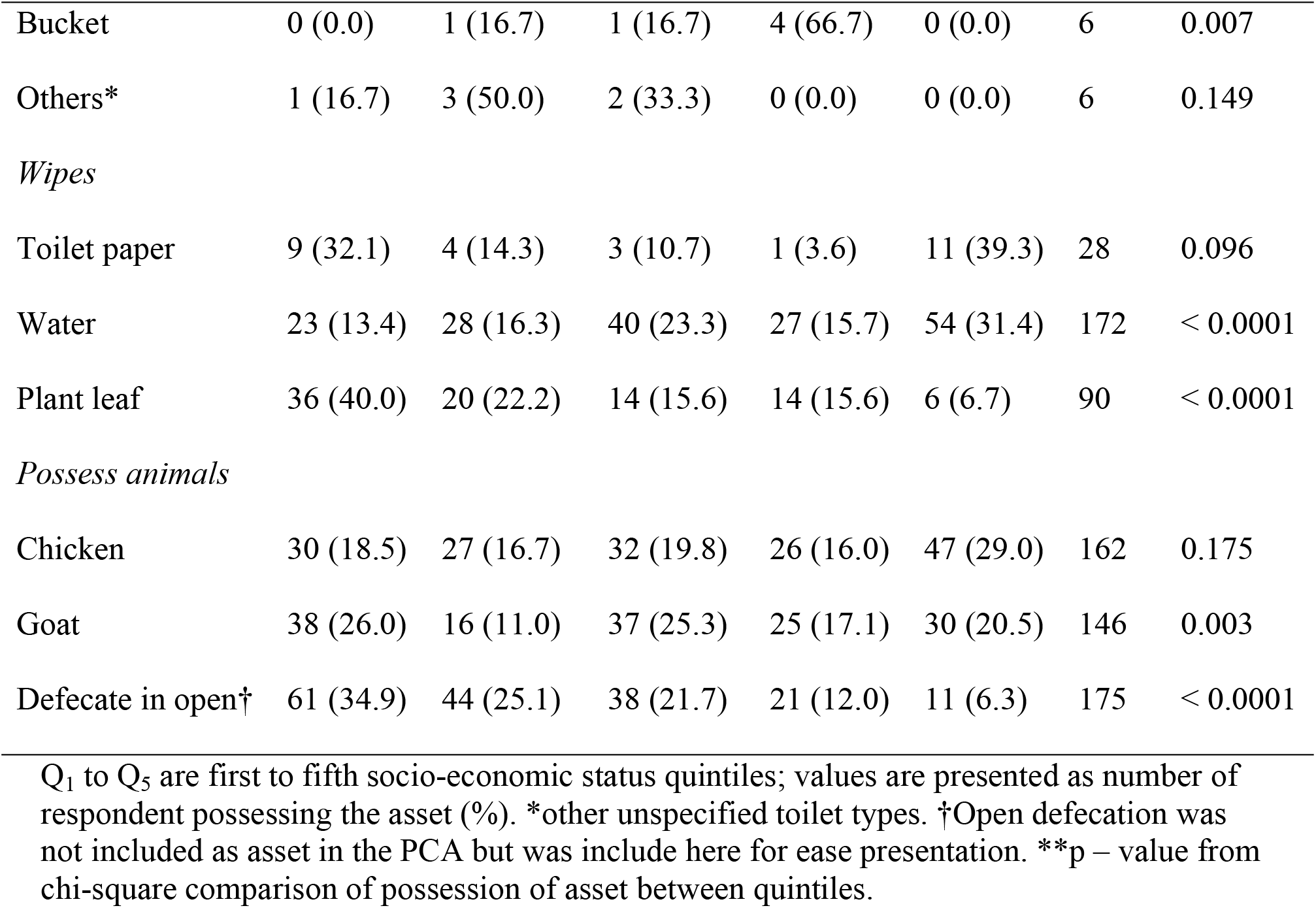
Wealth quintiles and relative possession of assets among agrarian communities in Ibaji and Igalamela-Odolu Local Government Areas, Kogi State.

| Descriptors | Wealth Quintiles (%) |  |  |  |  | Total | P** |
| --- | --- | --- | --- | --- | --- | --- | --- |
|  | Q<sub>1</sub> | Q<sub>2</sub> | Q<sub>3</sub> | Q<sub>4</sub> | Q<sub>5</sub> |  |  |
|  | (Poorest) | (Poor) | (Average) | (Rich) | (Richest) |  |  |
| **Total** | **68 (23.1)** | **52 (17.6)** | **58 (19.7)** | **44 (14.9)** | **73 (24.7)** | **295** |  |
| *Floor type* |  |  |  |  |  |  |  |
| Mud | 68 (45.6) | 49 (32.9) | 30 (20.1) | 2 (1.3) | 0 (0.0) | 149 | < 0.0001 |
| Cement | 0 (0.0) | 3 (2.1) | 28 (19.2) | 42 (28.8) | 73 (50.0) | 146 | < 0.0001 |
| *Electricity & Appliances* |  |  |  |  |  |  |  |
| Electricity | 2 (1.6) | 13 (10.6) | 17 (13.8) | 28 (22.8) | 63 (51.2) | 123 | < 0.0001 |
| Radio | 9 (5.3) | 28 (16.5) | 34 (20.0) | 29 (17.1) | 70 (41.2) | 170 | < 0.0001 |
| Television | 10 (12.8) | 6 (7.7) | 14 (17.9) | 17 (21.8) | 31 (39.7) | 78 | < 0.0001 |
| Refrigerators | 1 (4.8) | 0 (0.0) | 4 (19.0) | 6 (28.6) | 10 (47.6) | 21 | 0.005 |
| *Source of water* |  |  |  |  |  |  |  |
| Pipe-borne | 3 (2.0) | 16 (10.5) | 37 (24.2) | 27 (17.6) | 70 (45.8) | 153 | < 0.0001 |
| River | 67 (49.6) | 35 (25.9) | 18 (13.3) | 15 (11.1) | 0 (0.0) | 135 | < 0.0001 |
| Well | 1 (6.7) | 3 (20.0) | 3 (20.0) | 6 (40.0) | 2 (13.3) | 15 | 0.052 |
| Rain | 0 (0.0) | 0 (0.0) | 0 (0.0) | 7 (63.6) | 4 (36.4) | 11 | < 0.0001 |
| Have toilet | 2 (1.6) | 10 (7.8) | 24 (18.8) | 28 (21.9) | 64 (50.0) | 128 | < 0.0001 |
| *Toilet types* |  |  |  |  |  |  |  |
| Pit | 0 (0.0) | 5 (6.0) | 12 (14.3) | 17 (20.2) | 50 (59.5) | 84 | < 0.0001 |
| Water closet | 1 (3.1) | 0 (0.0) | 10 (31.2) | 7 (21.9) | 14 (43.8) | 32 | < 0.0001 |
| Bucket | 0 (0.0) | 1 (16.7) | 1 (16.7) | 4 (66.7) | 0 (0.0) | 6 | 0.007 |
| Others* | 1 (16.7) | 3 (50.0) | 2 (33.3) | 0 (0.0) | 0 (0.0) | 6 | 0.149 |
| *Wipes* |  |  |  |  |  |  |  |
| Toilet paper | 9 (32.1) | 4 (14.3) | 3 (10.7) | 1 (3.6) | 11 (39.3) | 28 | 0.096 |
| Water | 23 (13.4) | 28 (16.3) | 40 (23.3) | 27 (15.7) | 54 (31.4) | 172 | < 0.0001 |
| Plant leaf | 36 (40.0) | 20 (22.2) | 14 (15.6) | 14 (15.6) | 6 (6.7) | 90 | < 0.0001 |
| *Possess animals* |  |  |  |  |  |  |  |
| Chicken | 30 (18.5) | 27 (16.7) | 32 (19.8) | 26 (16.0) | 47 (29.0) | 162 | 0.175 |
| Goat | 38 (26.0) | 16 (11.0) | 37 (25.3) | 25 (17.1) | 30 (20.5) | 146 | 0.003 |
| Defecate in open† | 61 (34.9) | 44 (25.1) | 38 (21.7) | 21 (12.0) | 11 (6.3) | 175 | < 0.0001 |

Q<sub>1</sub> to Q<sub>5</sub> are first to fifth socio-economic status quintiles; values are presented as number of respondent possessing the asset (%). \*other unspecified toilet types. †Open defecation was not included as asset in the PCA but was include here for ease presentation. \*\*p – value from chi-square comparison of possession of asset between quintiles.

### Socioeconomic Status (SES) in Ibaji and Igalamela-Odolu LGAs

A total of 68 (23.1%) of the respondents were classified as poorest based on the 20-assets criteria. Those in richest category were 73 (24.7%) while poor, average and rich were slightly fewer (Table 4). Homes with mud floor was mainly associated with poverty, 68 (45.6%) and 49 (32.9%) of those that used mud floor were poorest and poor respectively. Majority of the people who had cement floors at home where in the fourth and fifth quintiles. There was very high significant difference in the distribution of floor types by wealth quintile (p < 0.0001). Electricity and electrical appliances such as radio, television and refrigerators were common in homes of the rich and richest members of the communities. Usage of toilet paper was not dependent on wealth quintiles. The poorest and poor were relatively more likely to practice open defecation and constituted 39.4% and 25.1% respectively of 175 who practiced it. This contrasts strongly with 12.0% and 6.3% of the rich and richest people respectively. Majority of people in Igalamela-Odolu LGA had better wealth status than those in Ibaji LGA (Fig 3).

**Fig 3.**
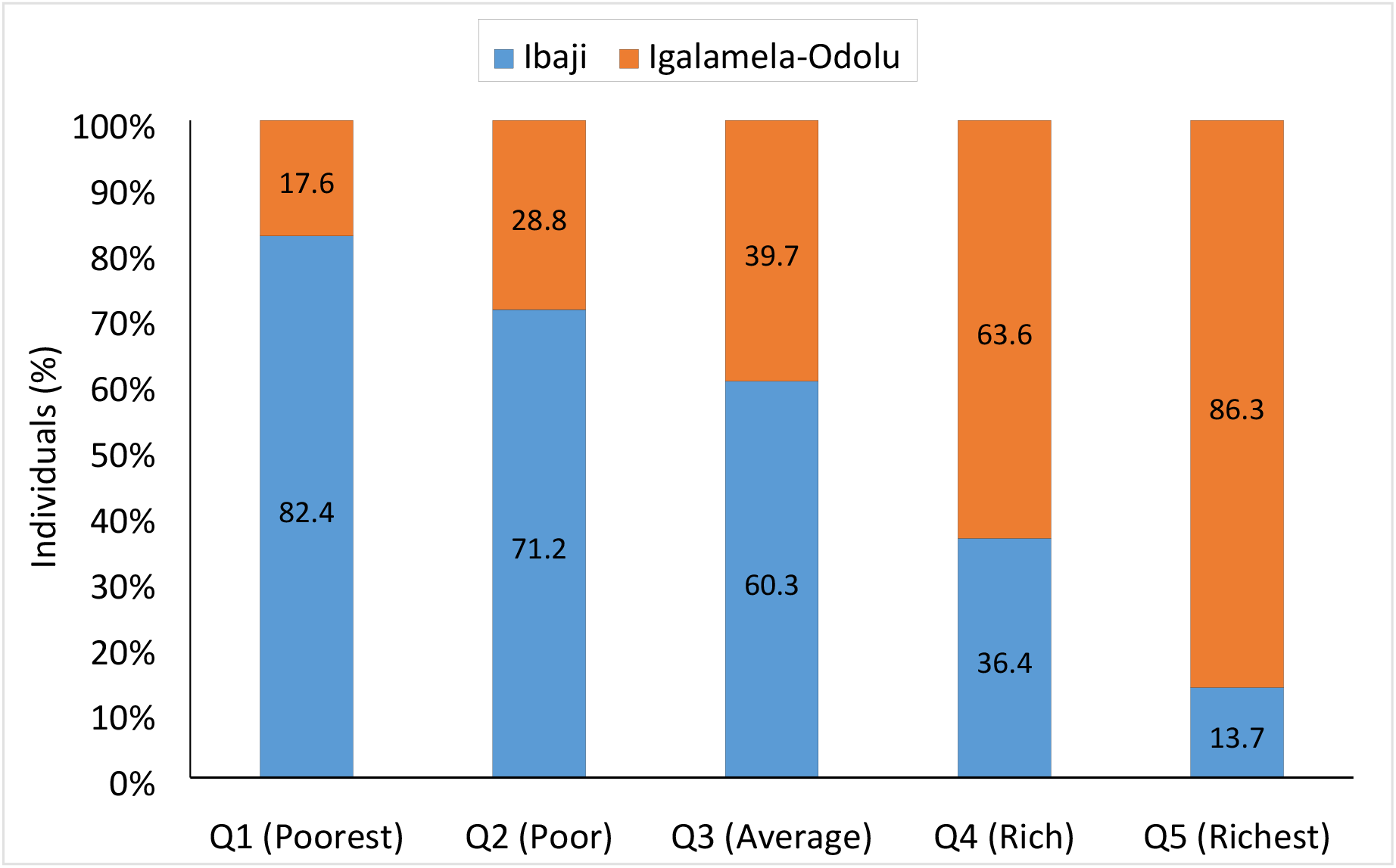
Distribution of wealth among individuals in Ibaji and Igalamela-Odolu Local Government Areas.

A summary of how STH infection was distributed in relation to the 20 assets is provided in supplementary file 2. Four assets were each significant determinant of STH infection. The single most important determinant was possession of toilet (χ^2^ = 13.469, p < 0.0001), especially pit toilet (χ^2^ = 10.222, p = 0.0001). The other two were ‘having water to wash after toilet’ (χ^2^ = 3.854, p = 0.050) and ‘using plant leaf as toilet paper’ (χ^2^ = 4.759, p = 0.029), and both more likely are associated with use of toilet and open defecation respectively. As was expected, people who had cement floor, electricity, television and tap water had lower cases of infection compared to those who did not (S2 Fig). Surprisingly, refrigerator as an indicator of wealth was associated with greater infection.

### Risk for infection with soil-transmitted helminths in Ibaji and Igalamela-Odolu LGAs

Risks of STH infection due to SES, knowledge, attitude and practices are presented in Tables 6 and 7. Risk of STH infection was similar between the first four wealth quintiles, poorest, poor, average and rich. However, risk of infection was significantly lower by 60% in the richest wealth quintile compared to the poorest quintile (PR = 0.4843, 95% CI = 0.2704 – 0.8678, p = 0.015). People that claimed to know parasitic worms, that think they could get worm, know how to get worm, and believed they had previously been infected by worms had little to no reduction in risk of STH infection when compared to those whose knowledge and perceptions were otherwise (Table 6). Open defecation was one major factor associated with significant increase in STH risk. People who practiced open defecation were two times more likely to harbor STH compared to those who did not (PR = 1.7878, 95% CI = 1.2366 – 2.5846, p = 0.00201). Compared to non-possession of toilet facilities, pit latrine and water closet toilet each reduced STH infection by approximately 50% (p < 0.05). Washing hands after toilet use, and wearing shoes while working in farms each also significantly lowered the risk of STH infection. Though washing of hands after toilet significantly lowered risk of infection, using soap for this purpose had no to little effect (Table 6).

**Table 6.**
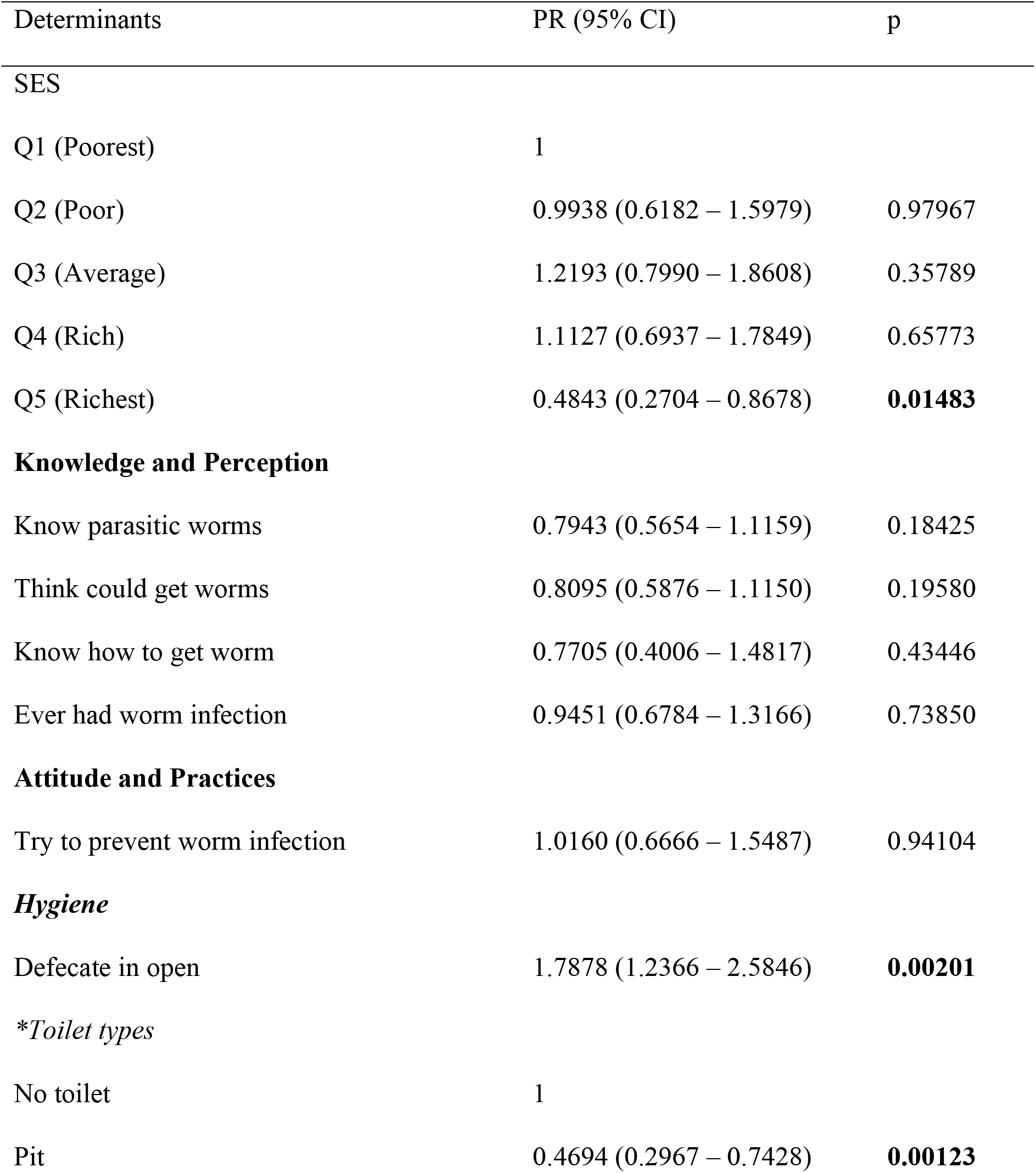
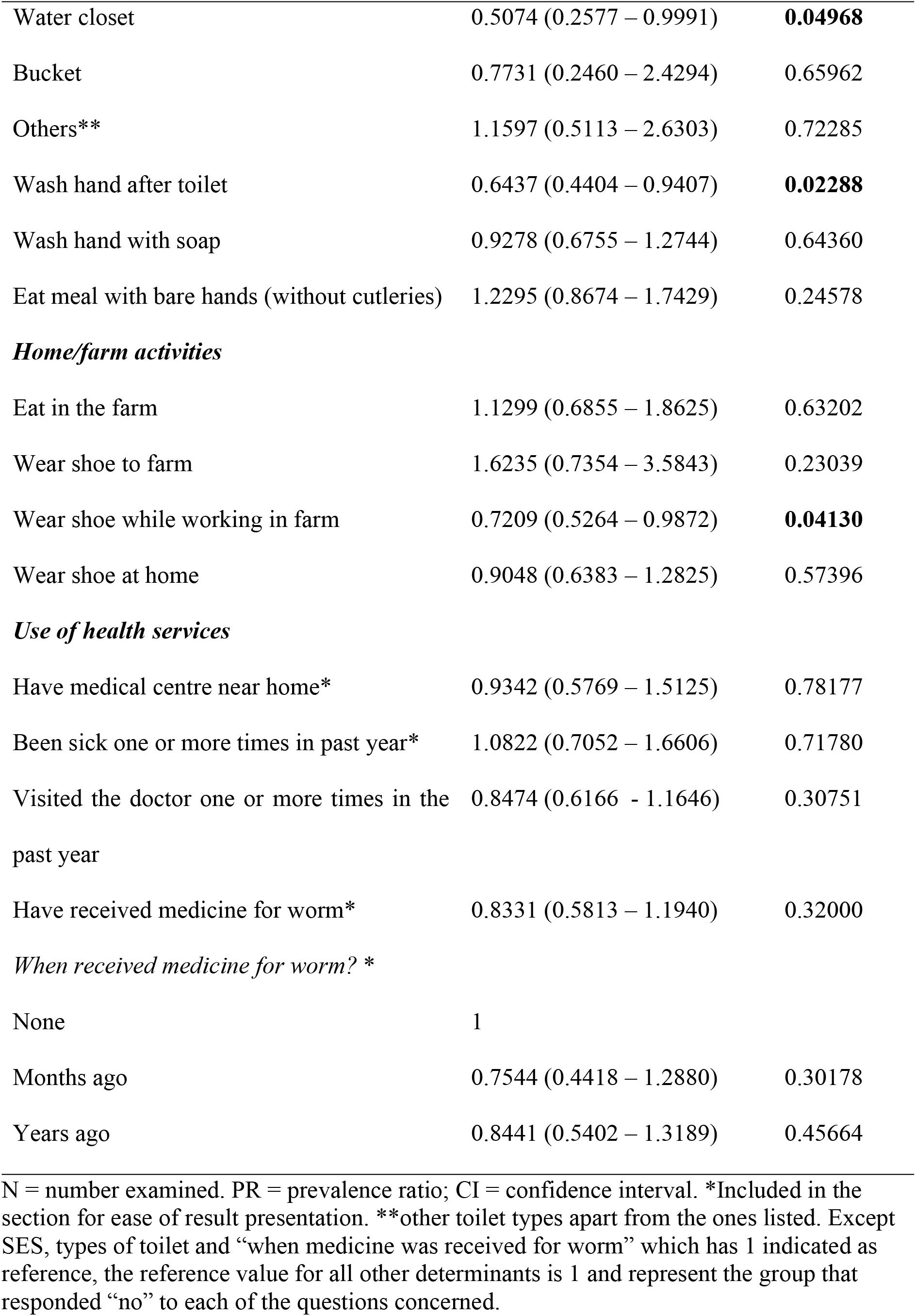
Risk of soil-transmitted helminths infection associated with socio-economic status (SES), knowledge, perception, attitude and practices among agrarian communities in Ibaji and Igalamela-Odolu Local Government Area, Kogi State. N = 295.

**Table 7.**
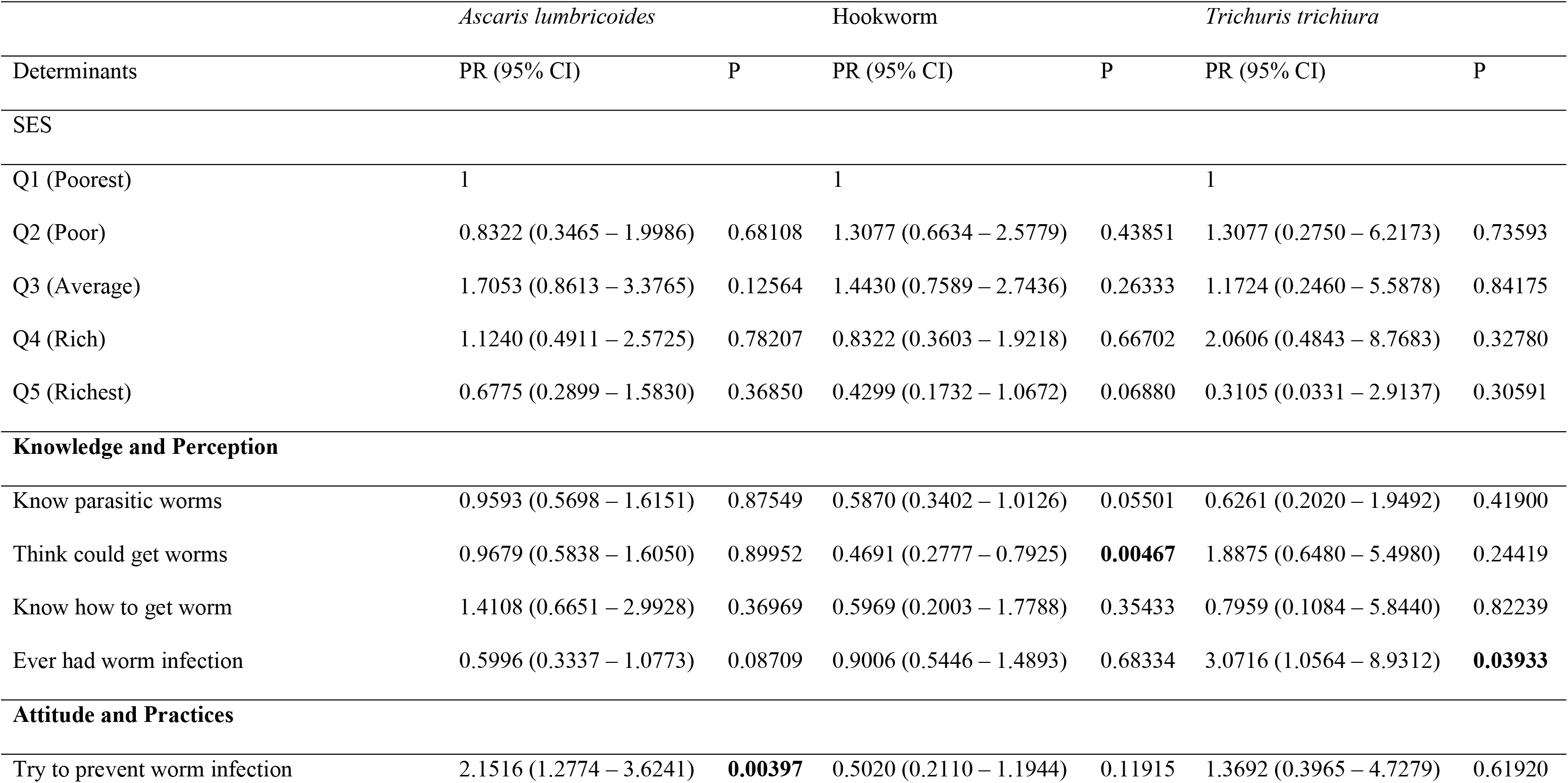
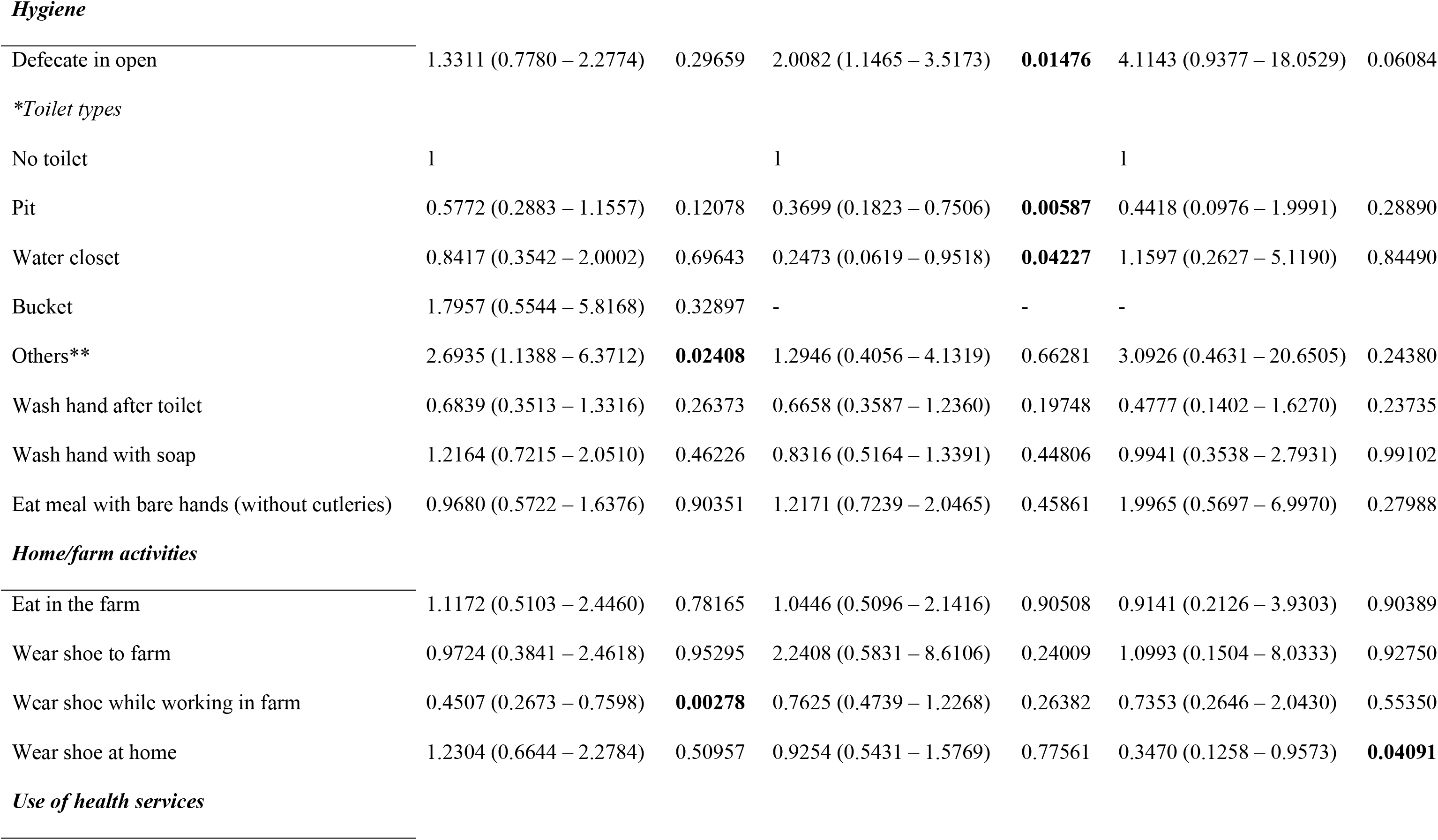
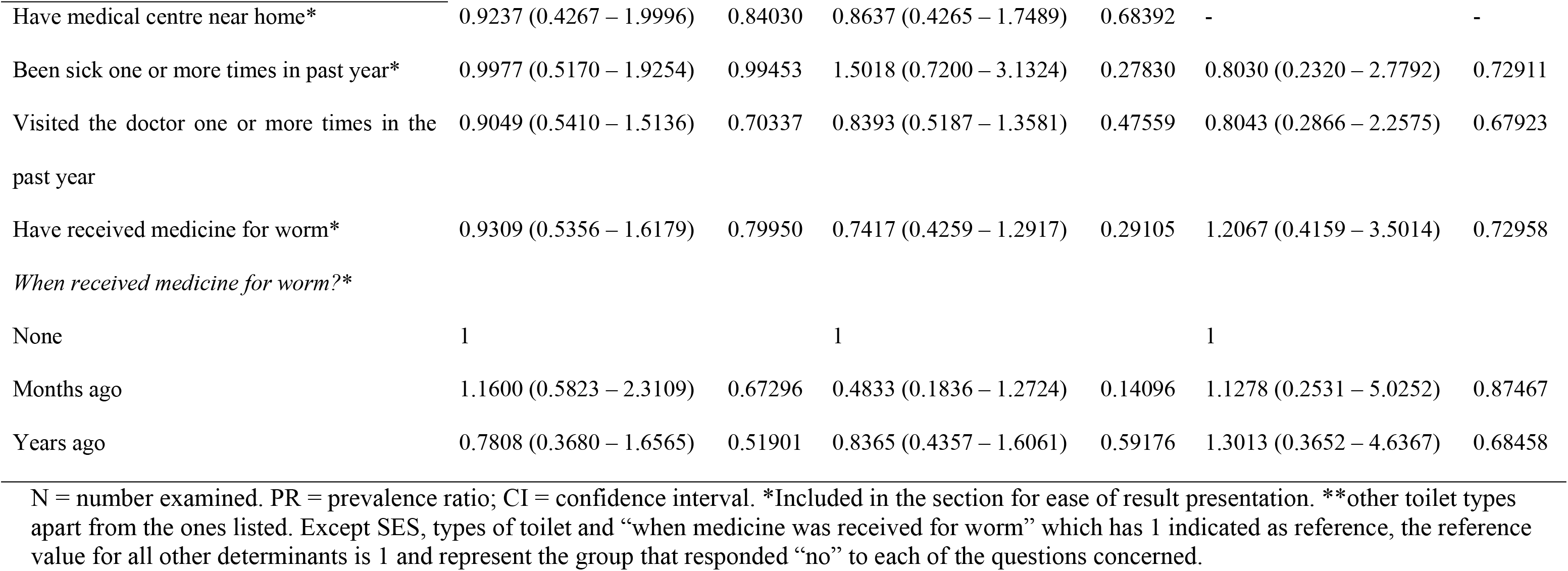
Risk of infection with *Ascaris lumbricoides*, hookworm and *Trichuris trichiura* associated with socio-economic status (SES), knowledge, attitude and practices (KAP) among agrarian communities in Ibaji and Igalamela-Odolu Local Government Area, Kogi State. (N = 295).

|  | *Ascaris lumbricoides* |  | Hookworm |  | *Trichuris trichiura* |  |
| --- | --- | --- | --- | --- | --- | --- |
| Determinants | PR (95% CI) | P | PR (95% CI) | P | PR (95% CI) | P |
| SES |  |  |  |  |  |  |
| Q1 (Poorest) | 1 |  | 1 |  | 1 |  |
| Q2 (Poor) | 0.8322 (0.3465 – 1.9986) | 0.68108 | 1.3077 (0.6634 – 2.5779) | 0.43851 | 1.3077 (0.2750 – 6.2173) | 0.73593 |
| Q3 (Average) | 1.7053 (0.8613 – 3.3765) | 0.12564 | 1.4430 (0.7589 – 2.7436) | 0.26333 | 1.1724 (0.2460 – 5.5878) | 0.84175 |
| Q4 (Rich) | 1.1240 (0.4911 – 2.5725) | 0.78207 | 0.8322 (0.3603 – 1.9218) | 0.66702 | 2.0606 (0.4843 – 8.7683) | 0.32780 |
| Q5 (Richest) | 0.6775 (0.2899 – 1.5830) | 0.36850 | 0.4299 (0.1732 – 1.0672) | 0.06880 | 0.3105 (0.0331 – 2.9137) | 0.30591 |
| **Knowledge and Perception** |  |  |  |  |  |  |
| Know parasitic worms | 0.9593 (0.5698 – 1.6151) | 0.87549 | 0.5870 (0.3402 – 1.0126) | 0.05501 | 0.6261 (0.2020 – 1.9492) | 0.41900 |
| Think could get worms | 0.9679 (0.5838 – 1.6050) | 0.89952 | 0.4691 (0.2777 – 0.7925) | **0.00467** | 1.8875 (0.6480 – 5.4980) | 0.24419 |
| Know how to get worm | 1.4108 (0.6651 – 2.9928) | 0.36969 | 0.5969 (0.2003 – 1.7788) | 0.35433 | 0.7959 (0.1084 – 5.8440) | 0.82239 |
| Ever had worm infection | 0.5996 (0.3337 – 1.0773) | 0.08709 | 0.9006 (0.5446 – 1.4893) | 0.68334 | 3.0716 (1.0564 – 8.9312) | **0.03933** |
| **Attitude and Practices** |  |  |  |  |  |  |
| Try to prevent worm infection | 2.1516 (1.2774 – 3.6241) | **0.00397** | 0.5020 (0.2110 – 1.1944) | 0.11915 | 1.3692 (0.3965 – 4.7279) | 0.61920 |
| ***Hygiene*** |  |  |  |  |  |  |
| Defecate in open | 1.3311 (0.7780 – 2.2774) | 0.29659 | 2.0082 (1.1465 – 3.5173) | **0.01476** | 4.1143 (0.9377 – 18.0529) | 0.06084 |
| *\*Toilet types* |  |  |  |  |  |  |
| No toilet | 1 |  | 1 |  | 1 |  |
| Pit | 0.5772 (0.2883 – 1.1557) | 0.12078 | 0.3699 (0.1823 – 0.7506) | **0.00587** | 0.4418 (0.0976 – 1.9991) | 0.28890 |
| Water closet | 0.8417 (0.3542 – 2.0002) | 0.69643 | 0.2473 (0.0619 – 0.9518) | **0.04227** | 1.1597 (0.2627 – 5.1190) | 0.84490 |
| Bucket | 1.7957 (0.5544 – 5.8168) | 0.32897 | - | - | - |  |
| Others** | 2.6935 (1.1388 – 6.3712) | **0.02408** | 1.2946 (0.4056 – 4.1319) | 0.66281 | 3.0926 (0.4631 – 20.6505) | 0.24380 |
| Wash hand after toilet | 0.6839 (0.3513 – 1.3316) | 0.26373 | 0.6658 (0.3587 – 1.2360) | 0.19748 | 0.4777 (0.1402 – 1.6270) | 0.23735 |
| Wash hand with soap | 1.2164 (0.7215 – 2.0510) | 0.46226 | 0.8316 (0.5164 – 1.3391) | 0.44806 | 0.9941 (0.3538 – 2.7931) | 0.99102 |
| Eat meal with bare hands (without cutleries) | 0.9680 (0.5722 – 1.6376) | 0.90351 | 1.2171 (0.7239 – 2.0465) | 0.45861 | 1.9965 (0.5697 – 6.9970) | 0.27988 |
| ***Home/farm activities*** |  |  |  |  |  |  |
| Eat in the farm | 1.1172 (0.5103 – 2.4460) | 0.78165 | 1.0446 (0.5096 – 2.1416) | 0.90508 | 0.9141 (0.2126 – 3.9303) | 0.90389 |
| Wear shoe to farm | 0.9724 (0.3841 – 2.4618) | 0.95295 | 2.2408 (0.5831 – 8.6106) | 0.24009 | 1.0993 (0.1504 – 8.0333) | 0.92750 |
| Wear shoe while working in farm | 0.4507 (0.2673 – 0.7598) | **0.00278** | 0.7625 (0.4739 – 1.2268) | 0.26382 | 0.7353 (0.2646 – 2.0430) | 0.55350 |
| Wear shoe at home | 1.2304 (0.6644 – 2.2784) | 0.50957 | 0.9254 (0.5431 – 1.5769) | 0.77561 | 0.3470 (0.1258 – 0.9573) | **0.04091** |
| ***Use of health services*** |  |  |  |  |  |  |
| Have medical centre near home* | 0.9237 (0.4267 – 1.9996) | 0.84030 | 0.8637 (0.4265 – 1.7489) | 0.68392 | - | - |
| Been sick one or more times in past year* | 0.9977 (0.5170 – 1.9254) | 0.99453 | 1.5018 (0.7200 – 3.1324) | 0.27830 | 0.8030 (0.2320 – 2.7792) | 0.72911 |
| Visited the doctor one or more times in the past year | 0.9049 (0.5410 – 1.5136) | 0.70337 | 0.8393 (0.5187 – 1.3581) | 0.47559 | 0.8043 (0.2866 – 2.2575) | 0.67923 |
| Have received medicine for worm* | 0.9309 (0.5356 – 1.6179) | 0.79950 | 0.7417 (0.4259 – 1.2917) | 0.29105 | 1.2067 (0.4159 – 3.5014) | 0.72958 |
| *When received medicine for worm?\** |  |  |  |  |  |  |
| None | 1 |  | 1 |  | 1 |  |
| Months ago | 1.1600 (0.5823 – 2.3109) | 0.67296 | 0.4833 (0.1836 – 1.2724) | 0.14096 | 1.1278 (0.2531 – 5.0252) | 0.87467 |
| Years ago | 0.7808 (0.3680 – 1.6565) | 0.51901 | 0.8365 (0.4357 – 1.6061) | 0.59176 | 1.3013 (0.3652 – 4.6367) | 0.68458 |

N = number examined. PR = prevalence ratio; CI = confidence interval. \*Included in the section for ease of result presentation. \*\*other toilet types apart from the ones listed. Except SES, types of toilet and “when medicine was received for worm” which has 1 indicated as reference, the reference value for all other determinants is 1 and represent the group that responded “no” to each of the questions concerned.

Risk of *A. lumbricoides* infection was similar between the first four quintiles. Though people in the fifth wealth quintiles had about 40% reduced risk compared to first quintile, but this was not significant (p = 0.36850). People who responded affirmatively to making effort at preventing worm infection unfortunately had more than 2 times greater risk of *A. lumbricoides* infection (Table 6). The greater risk among those who “try to prevent worm infection” compared to those who did not ranged from 30% to 360%, and was significant (p = 0.00397). Wearing shoes while working in farms also significantly lowered the risk of *A. lumbricoides* infection by over 50% (p = 0.00278). Those that practiced open defecation were at 30% greater risk of *A. lumbricoides* infection as those who did not (Table 7).

Risk of hookworm infection dropped in the fourth and fifth wealth quintiles compared to the first (Table 7). The drop was more obvious among the richest people, though it was not significant (p = 0.06880). Those who were conscious of the possibility of getting infected by worms had lower chances of infection with hookworm; risk of infection dropped significantly by over 50% (p = 0.00467). Open defecation increased risk of hookworm infection by 200% (PR = 2.0082, 95% CI = 1.1465 – 3.5173, p = 0.01476). Pit latrine and water closet toilets compared to non-toilet owners reduced risk of infection by 65% and 70% respectively, the differences were significant (p < 0.05).

People who acknowledged being previously infected by worm were at over 300% greater risk of *T. trichiura* infection. The actual range of this estimate was 5% to 800%, and was significant (PR = 3.0716, 95% CI = 1.0564 – 8.9312, p = 0.03933). Open defecation also increased the risk of infection by about 400% though this was not significant (p > 0.05; Table 7). Those that wore shoes at home were 70% less likely to get infected by *T. trichiura* (p = 0.04091).

## Discussion

Overall prevalence of STH was 34.2%. STH isolated were *Ascaris lumbricoides*, hookworm and *Trichuris trichiura*, and at prevalence 18.1%, 16.8% and 5.2% respectively. Intensities of the three helminths were largely light. Collectively, prevalence of moderate and heavy intensity of STH infection was < 1%. A total of 68 (23.1%) of the respondents were classified as poorest based on the 20-assets criteria. Those in richest category were 73 (24.7%) while poor, average and rich were slightly fewer. Mud floor at homes was mainly associated with poverty. Majority of the people who had cement floors at home where in the fourth (rich) and fifth (richest) quintiles. There was very high significant difference in the distribution of mud and cement floors by wealth quintile (p < 0.0001). The poorest and poor were relatively more likely to practice open defecation and constituted 39.4% and 25.1% respectively of 175 who practiced it; which contrasts strongly with 12.0% and 6.3% of the rich and richest people respectively. Majority of people in Igalamela-Odolu LGA had better wealth status than those in Ibaji LGA. Risk of infection was significantly lower by 60% in the richest wealth quintile compared to the poorest quintile. People who practiced open defecation were two times more likely to harbor STH compared to those who did not. Compared to non-possession of toilet facilities, pit latrine and water closet toilet each significantly reduced the risk STH infection. Washing hands after toilet use significantly lowered the risk of STH infection; though using soap for this purpose had no to little effect.

Overall prevalence of STH in Ibaji and Igalamela-Odolu LGAs was moderate (34.2%). This falls within the 16.2% – 39.0% range among LGAs of Kogi State as earlier reported by the Nigerian Federal Ministry of Health [18]. However, FMOH survey was only school-based. Most STH prevalence surveys in Nigeria were carried out among school children: Sowemino and Asaolu [31] obtained prevalence of 34.4% in Ile-Ife, Osun State; 26.66% was reported in Rivers State [32], 80.9% among Almajiri children in northern Nigeria [33], and 30.3% in Imo state [9]. Differences in prevalence obtained in various parts of the country relative to the present study may be attributable to environmental, ecological and anthropogenic factors that prevailed in each area. Moreover, methodological differences between studies could also be an issue; while floatation techniques were applied in most of these surveys, the standard Kato-Katz technique was employed in this study. Comparative studies on floatation and Kato-Katz techniques have reported both similar and disparate performance depending on the helminth of interest [34–36]. Despite the 34.2% prevalence, mean intensity of STH was low; moderate to high intensity was below 1% in the study population. This finding holds a promise in terms of possibilities of control of STH in the population.

An important aspect of the result is the similarity in infection among the four age groups (preschool, school, women of reproductive age and men of same age, and older at-risk groups). This further emphasizes need for extension of preventive chemotherapy to other at-risk groups, especially in settings such as the one covered by this study where majority of the inhabitants are at risk. The finding corroborates STHAC recommendation [16,17]. Though women of reproductive age were considered along with men of similar age group (52.8% vs. 47.2%), there are no bases for expecting that helminth burden will be any different between men and women considering the locality in question. Infection was also not significantly different between male and female (unreported). Restricting mass drug administration (MDA) to preschool-aged and school-aged in agrarian communities of Ibaji and Igalamela-Odolu LGAs, and other localities with similar attributes only increases the risk of re-infection after MDA.

Factors associated with STH infection were identified in this study. Socio-economic and demographic factors such as poverty, lack of portable water, occupation, age, and attitudes such as poor sanitation and poor sewage disposal have been recognized as determinants of STH distribution in endemic areas [15,37]. From the asset-based approach used in this study, the number of very poor and very rich households was almost equal in the study area. While the asset-based approach employed effectively classified the household into wealth quintiles, number and types of assets considered generally influence classifications [27]. From the present study, people at the fifth wealth quintile (richest) were less likely to harbour STH. Possession of assets indicative of good SES such as cement floor, electricity, radio, television, pipe-borne water and toilet facilities such as pit latrine and water closets reduced risk of STH. Open defecation which was common among people in the first, second and third wealth quintile was a major risk for STH infection. It has been stated previously that improper sewage disposal is a major factor for STH infection [38]. Availability of toilet facilities in a household is a matter of social class proving the saying that poverty breeds diseases. Mud and hut house is associated with increased STH infection [39]. Mud has the tendency of retaining parasite eggs even when it has been apparently cleaned. Mud floor which was assigned strong negative factor score by statistical analysis is one of the strong indicators of poverty in the LGAs. In addition, cement floor alone was one of the SES assets that strongly reduced the risk of infection with STH. Thus, STH infection in the LGAs can largely be blamed on SES. Though it has been suggested by Becker et al. [17], that it is unlikely that significant impact can be made between 2018 and 2020 concerning socio-economic status of people in STH endemic areas. This therefore suggests, that other approaches such as preventive chemotherapy are more important if the WHO 2020 target is to be achieved. The authors share the same view, but would wish to emphasize that in order to sustain gains, a sustainable approach against STH requires improved SES.

Knowledge, attitude and practices were other strong determinants of STH infection in the LGAs. Fifty-nine percent (59%) of the respondents admitted defecating in the open mainly due to lack of toilets. Risk of STH infection was generally higher among those who practiced open defecation. This was especially obvious for hookworm and *T. trichiuria*. This corroborates reports from Nigeria and other parts of the world [39,40]. Regular visits to defecation spots increase chances of hookworm infection because the infective stages of this parasites develop around defecation sites where the eggs were released and are always ready to strike and infect an unsuspecting host. Open defecation can be reduced through health education and by improving the SES of communities affected through provision of amenities such as pit latrines and water closet toilets.

Hand washing after toilet use lowered the risk of acquiring STH. Although auto-infection from freshly passed stool is not possible since period of incubation in the soil is required by STH, habitual hand washing after toilet use is an indication of high sense of personal hygiene. More so, individuals, especially children, who use regular open defecation sites are more likely to soil their hands with previously contaminated soil. Therefore hand washing protects against STH. It has been advocated that health education in the aspect of hand washing with soap be intensified among communities where STH infection occurs [41]. Though the present study found no special role played by soap. The importance of water, sanitation and hygiene (WASH) which was re-affirmed by STHAC is emphasized by the present study.

Chances of having STH infection were also lowered significantly by wearing shoes while working on the farm. This act provided some level of protection against the three STH, and the effect was significant against *A. lumbricoides*. Not wearing of shoes was similarly reported as a risk factor for hookworm infection in northwestern Ethiopia [42]. The infective hookworm larvae from soil enter into the host by active penetration of unbroken skin especially the feet that are often in contact with the soil. Shoe wearing therefore, reduces the risk of hookworm infection. Though unrelated to *A. lumbricoides* infection, shoe wearing prevents contamination of feet which could indirectly lead to infection when individuals touch such contaminated feet and eat with unwashed hands afterwards.

### Conclusion and recommendations

In conclusion, this study reports a moderate prevalence and light intensity of STH infection in the study population indicating its endemicity in the study area and therefore a public health problem that requires attention. A number of risk factors associated with STH transmission was identified. These are mainly socio-economic, attitude and practices. Based on the findings, it is hereby recommended that

1. Mass preventive chemotherapy against STH infection through the interventions of the Ministry of Health and other health organizations should be extended to all age groups in the affected communities. This is especially important to prevent risk of re-infection by other at-risk adults.
2. Health education and awareness on transmission and health impact of STH should be emphasized. The study further affirms water, sanitation and hygiene (WASH) as integral STH control strategy.
3. Effort should be made at improving SES in places where STH has moderate to high endemicity such as Ibaji and Igalamela-Odolu LGAs through provision of basic amenities such as pit latrine and water closet toilets, pipe-borne water, electricity and good housing.

## Acknowledgements

We thank the traditional heads in Ibaji and Igalamela-Odolu LGAs, and the households that willing participated in the study.

## Supporting information

**S1 Table. Supporting information from the asset-based socio-economic status**

**S2 Fig. Proportion of people positive and negative for soil-transmitted helminthiasis who had different assets.** The five key determinants are highlighted in dark-walled rectangles. Possession of toilet was the most important determinant, usage of plant leaf or water is closely associated with possession of toilet (open defecation is not an asset, but was included here for convenience of presentation).

**S3 Checklist: STROBE Checklist**

